# Multiple evolutionary trajectories for non-O157 Shiga toxigenic *Escherichia coli*

**DOI:** 10.1101/549998

**Authors:** Nabil-Fareed Alikhan, Nathan L. Bachmann, Nouri L. Ben Zakour, Nicola K. Petty, Mitchell Stanton-Cook, Jayde A. Gawthorne, Donna M. Easton, Timothy J. Mahony, Rowland Cobbold, Mark A. Schembri, Scott A. Beatson

## Abstract

**Background:** Shiga toxigenic *Escherichia coli* (STEC) is an emerging global pathogen and remains a major cause of food-borne illness with more severe symptoms including hemorrhagic colitis and hemolytic-uremic syndrome. Since the characterization of the archetypal STEC serotype, *E. coli* O157:H7, more than 250 STEC serotypes have been defined. Many of these non-O157 STEC are associated with clinical cases of equal severity as O157. In this study, we utilize whole genome sequencing of 44 STEC strains from eight serogroups associated with human infection to establish their evolutionary relationships and contrast this with their virulence gene profiles and established typing methods.

**Results:** Our phylogenomic analysis delineated these STEC strains into seven distinct lineages, each with a characteristic repertoire of virulence factors. Some lineages included commensal or other *E. coli* pathotypes. Multiple independent acquisitions of the Locus for Enterocyte Effacement were identified, each associated with a distinct repertoire of effector genes. Lineages were inconsistent with O-antigen typing in several instances, consistent with lateral gene transfer within the O-antigen locus. STEC lineages could be defined by the conservation of clustered regularly interspaced short palindromic repeats (CRISPRs), however, no CRISPR profile could differentiate STEC from other *E. coli* strains. Six genomic regions (ranging from 500 bp - 10 kbp) were found to be conserved across all STEC in this dataset and may dictate interactions with Stx phage lysogeny.

**Conclusions:** The genomic analyses reported here present non-O157 STEC as a diverse group of pathogenic *E. coli* emerging from multiple lineages that independently acquired mobile genetic elements that promote pathogenesis.

## Background

Food-borne pathogens persist as a major cause of clinical infection within the industrialized world (1,2). Shiga toxigenic *E. coli* (STEC) is one such emerging global food-borne pathogen responsible for severe human disease with symptoms of hemorrhagic colitis (HC) and hemolytic-uremic syndrome (HUS) (3,4).

Enterohemorrhagic *E. coli* (EHEC), a subset of STEC associated with human disease, were first identified in 1982 from an outbreak of contaminated beef (5). Since then, the serotype associated with that outbreak, O157:H7, has been linked to many other major outbreaks across the globe. The features used to serotype *E. coli* include the O-antigen of the lipopolysaccharide (LPS) and the H-antigen of the flagella (6). More than 150 different O antigens have been recognized and each one defines a specific serogroup (7). A combination of the O and H antigens is used to define a serotype (Nataro and Kaper, 1998). An additional 250 STEC serotypes have also been associated with human disease, often referred to collectively as non-O157 (8).

STEC are a heterogeneous group of *E. coli* that exhibit a high degree of genomic and phenotypic diversity. Only certain Shiga toxin harboring serogroups have been associated with human disease (i.e. are EHEC), and within these serogroups there are also differences in their association with human disease outbreaks (8,9). In the United States, these differences have been formalized into a ‘Seropathotyping’ scheme, which ranks serogroups by degree of pathogenesis based on outbreak prevalence and epidemiological studies. This scale places O157:H7 as the most prevalent STEC serotype followed by O26, O111, O103, O121 and O145, and finally O91, O104 and O113 (10). As yet, it is unclear how much of the variation observed between serogroups is linked to differences in the host response to infection, or is due to variations in virulence gene content between strains.

STEC harbor Shiga toxins (Stx), which are lambdoid phage-encoded verocytotoxins that are homologous to *Shigella dysenteriae* type 1 toxins and cause HUS in humans (5,6). Stx has been classified into two main groups (Stx1 and Stx2), with Stx1 and Stx2 both further divided into a number of more closely related subtypes (7). Common Stx variants have been summarized in Supplementary Table 1.

Other than encoding at least one *stx* gene, some STEC, including O157, also carry the Locus of Enterocyte Effacement (LEE) pathogenicity island, which encodes a Type III Secretion System (T3SS), the adhesion Intimin and its translocated receptor Tir (11,12). Many of the genes encoded within the LEE pathogenicity island are responsible for the attaching and effacing phenotype induced by EHEC and Enteropathogenic *E. coli* (EPEC) strains (13). LEE encodes the structural genes of the *E. coli* T3SS and up to six effector proteins (11). In addition, numerous other effector genes have been found scattered throughout the chromosome, many in clusters referred to as ‘exchangeable effector loci’ (EELs) and they are often associated with mobile genetic elements such as prophages (18).

A number of LEE-negative STEC are also associated with human infection. These strains have other genes that contribute to virulence, including *epeA, sab, and subAB* (14–16), and appear to adhere to epithelial cells via alternative mechanisms. Furthermore, a number of plasmid-encoded virulence genes have been identified in STEC, including those encoding hemolysin (*hlyA*), proteases (*espP* and *katP*) and cytotoxins (*subA*) (15,17,18). Despite extensive screening, no virulence factor or marker, other than the *stx* genes themselves, are universally conserved across all STEC although the combination of *stx_2_* and *eae* is associated with more severe clinical outcomes associated with EHEC infection (19).

STEC strains also exhibit diversity in their phylogeny, and are represented across the *E. coli* species. The O157 serotype is found within *E. coli* phylogroup E, having evolved from an ancestral O55:H7 EPEC strain (20). In contrast, the dominant non-O157 STEC serotypes are generally found within the B1 phylogroup (21). It has been established that non-O157 STEC, like other pathogenic *E. coli*, are comprised of multiple independent lineages that acquired key virulence factors through stepwise evolution (22–24). This concept of parallel evolution was first presented through phylogenetic analysis of seven housekeeping genes (25) and confirmed by whole genome sequencing of O26, O103 and O111 STEC strains (26). These studies demonstrated that STEC pathogenicity is derived from the acquisition of virulence factors via multiple independent events, and mediated via the acquisition of mobile genetic elements.

To establish the evolutionary relationship of non-O157 STEC, and provide a phylogenomic framework for investigating differences in virulence gene profile and established typing methods, we performed whole genome sequencing of forty-four genetically diverse STEC strains from eight serotypes commonly associated with human disease (O26, O111, O91, O128, O103, O113, O121 and O45). The strains were obtained from Australia and the United States, with the majority of strains representing the more clinically relevant serotypes O26 and O111. Overall, this study offers an overview of non-O157 STEC, including both LEE-negative and LEE-positive strains of clinical significance, presenting STEC as a diverse group of *E. coli* from at least seven distinct lineages with varying virulence profiles. Through the high granularity granted by next-generation sequencing, we were able to contrast STEC phylogeny and established typing methods, including O-antigen typing and EcMLST, to test whether these approaches prove valid against a whole genome phylogeny. We were also able to determine whether a previously published CRISPR typing method for LEE-positive non-O157 strains is applicable to STEC at large.

## Results

### Genome assembly

Forty-four STEC strains sourced from Australia and the United States, and originating from human, ruminant livestock and contaminated food origins, were investigated in this study (Additional File 1). These strains represented 20 different serotypes from serogroups associated with human disease including O26, O111, O91, O128, O103, O113, O121 and O45. The average read coverage for each genome for the 44 STEC strains was 252±134 times the total genome size. Each genome assembly had an average total length of 5,389,684±239,389 bp, and an average N50 scaffold size of 90,958 ±30,439 bp. The number of scaffolds within the assemblies ranged from 121 to 979 scaffolds (mean of 291).

### Phylogenetic analysis

To obtain a high-resolution overview of the non-O157 STEC genomes, a maximum-likelihood tree based on 2,153 aligned core gene sequences (including 48,912 variable sites) was generated for the 44 strains, as well as seven previously published *E. coli* reference genomes from the B1 phylogroup and *E. coli* O157 Sakai as an outgroup (Figure 1; Supplementary Table 2, Additional File 1). To ensure that the lineages defined in this study were not the product of rapid evolution due to recombination we also determined a recombination-free tree in which core genes were removed if there was significant evidence for homoplasy (Supplementary Figure 1). The recombination-free tree included 1,136 genes and exhibited the same phylogenetic topology for major STEC lineages (ST106A, ST106B, ST118, ST379,ST234, ST89 and ST461) as observed in Figure 1, with branches displaying >90% bootstrap support. The phylogenetic analysis revealed that STEC strains form multiple lineages that have evolved in parallel and are distinct from O157. Individual lineages could be classified according to a particular sequence type according to the EcMLST seven allele scheme (Figure 1 and Additional File 1).

**Figure 1.**
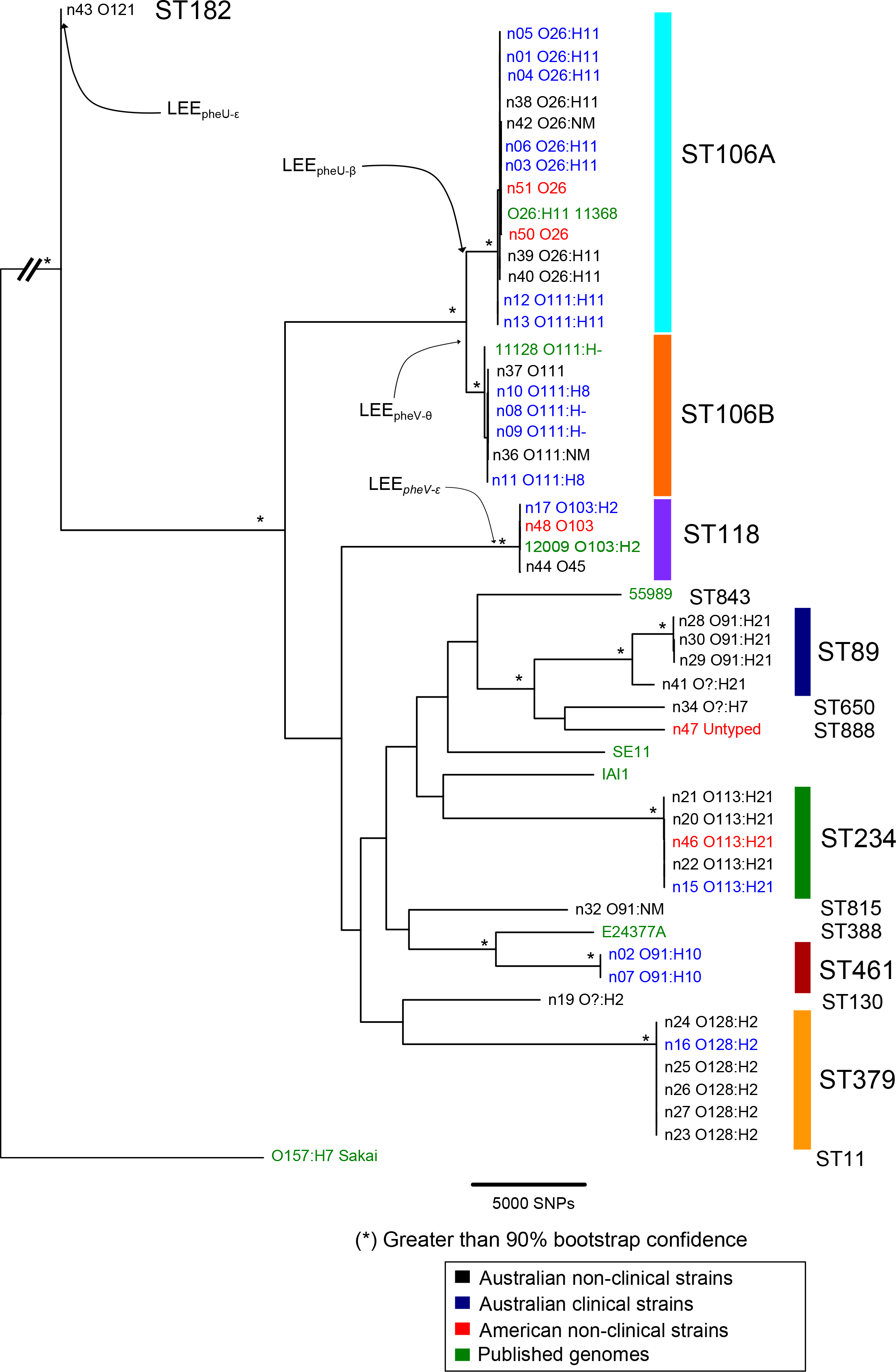
Phylogenetic relationship of non-O157 STEC with *E. coli* B1 Phylogroup. Maximum Likelihood (ML) phylogram with asterisks indicating bootstrap support greater than 90% from 400 replicates. The tree was rooted using *E. coli* O157:H7 Sakai. The phylogram includes forty-four Shiga toxin positive *E. coli* from this study and seven other *E. coli* strains from the B1 phylogroup including 55989, SE11, IAI1, E24377A, and previously sequenced non-O157 strains; 11368, 11128 and 12009. Accession numbers of these strains are listed in (Supplementary Table 2). Genomes have been annotated and highlighted according to lineage and named according to the EcMLST seven allele schema. Isolate sources are indicated in the key. The phylogram was built from 48,912 nucleotide SNPs from 2,153 *E. coli* genes, which are the number of genes conserved across the B1 Phylogroup, using PhyML (v20120412) (69) with the HKY85 substitution model. SplitsTree4 (82) was used to generate the final consensus tree. The final figure was prepared in FigTree (v1.4) (83). Labels indicate independent acquisition of the Locus for Enterocyte Effacement (LEE), site of insertion (*pheV*, *pheU*) and Intimin type (epsilon, theta and beta).

To enable a scalable nomenclature, we refer to clades by their EcMLST sequence types. Phylogroup B1 STEC strains formed a clade structure that was polyphyletic and intermingled with other non-STEC *E. coli* from the B1 phylogroup including EAEC, ETEC and commensal strains. Notably, ST461 strains were most closely related to ETEC strain E24377A rather than other STEC strains, indicating a common ancestor. Whereas, all O26 and O111 strains were ST106, but fell into two distinct sub-lineages that are referred to here as ST106A and ST106B. Of the 44 STEC strains sequenced in this study, three strains, n19, n32 (O91:NM) and n43 (O121), did not cluster with any other strains. Strain n43 (ST182) in particular was highly divergent to all B1 strains (Figure 1). In general, STEC clades defined in this study included a mixture of strains isolated from clinical samples, contaminated food or ruminant livestock.

### Shiga Toxins and the Locus of Enterocyte Effacement

Shiga toxin type was heterogeneous across strains within this study suggesting that the acquisition and loss of *stx* genes is dynamic and has occurred in multiple independent events. An explanation of the different Stx types can be found in Supplementary Table 1. Stx type (summarized in Figure 2 and Additional File 1) was consistent with several of the defined lineages, including ST106A, ST106B, ST461 and ST379. ST461 and ST379 strains carried *stx2d*, as well as *stx1c* and *stx2d_act_*, respectively. ST106A strains differed from ST106B strains; whereas ST106B strains carried both *stx1a* and *stx2a*, ST106A strains only carried *stx1a*. This suggests that *stx2a* was gained in ST106B, or alternately, was lost in ST106A after the two lineages diverged. There was also evidence for variation in *stx* content within the defined lineages examined. For example, n30 (ST89) contains *stx2d_act_* and *stx1a* while all other strains within this lineage only contain *stx2d_act_*. Similarly, strains n20 and n21 (ST234) were *stx2a* and *stx2d* positive. However, other ST234 strains contained either *stx2a* or *stx2d* (but not both), with acquisition of two *stx* prophages within the lineage followed by deletion within individual strains being the most parsimonious explanation. Due to the repetitive nature of the Stx encoding prophages, Stx genes did not assemble along with their cognate phage or the insertion site. Attempts to resolve this through mapping the underlying reads showed no unambiguous link between prophage and the Stx genes. As such, it was not possible to determine the Stx-encoding phage insertion site. This could be addressed using long-read sequencing technologies.

**Figure 2.**
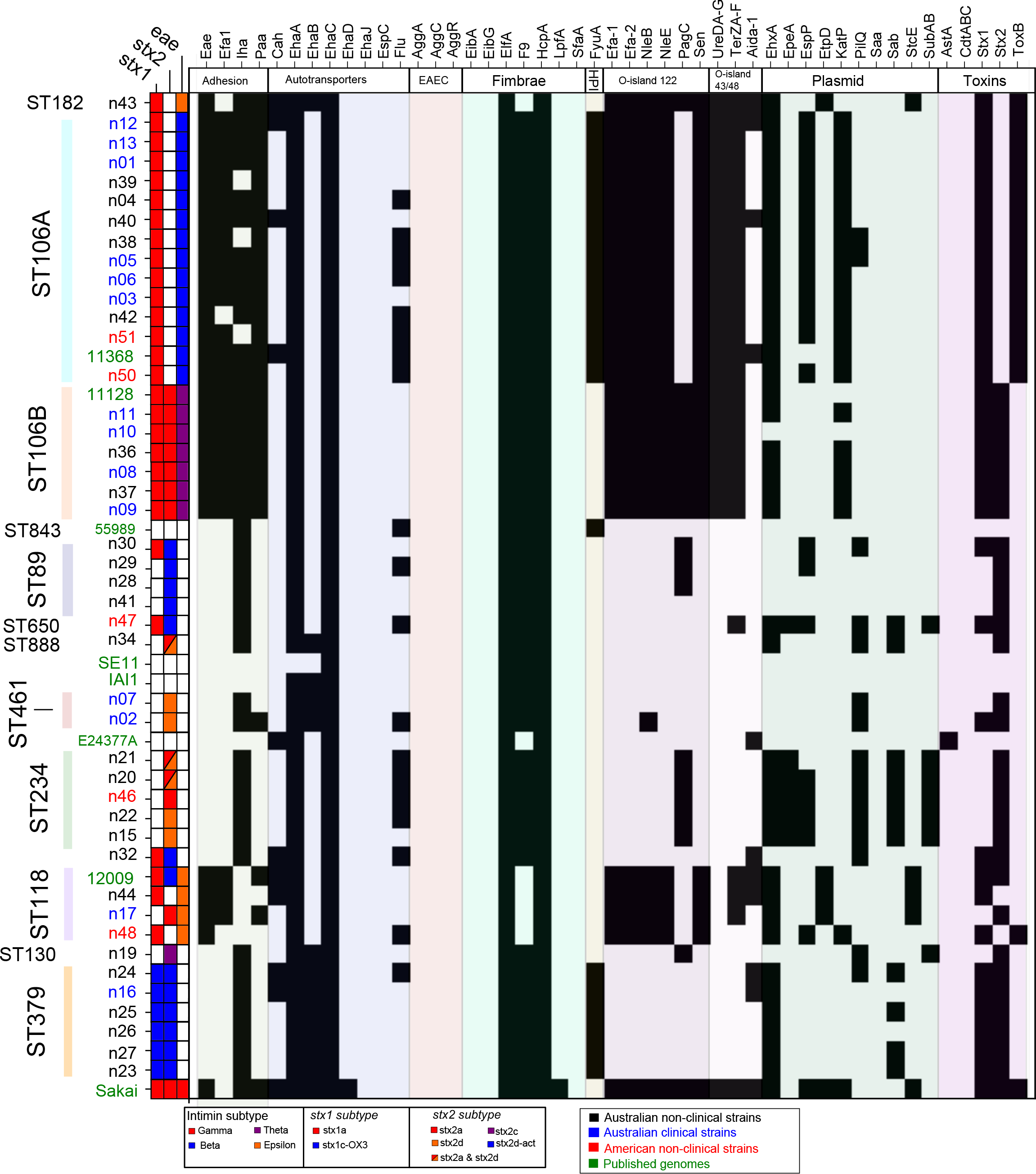
Virulence profile of non-O157 STEC and *E. coli* within B1 Phylogroup. Presence/absence matrix of a panel of STEC virulence factors. Virulence factors are shown along the x-axis with strains along the y-axis, listed in the order presented in the whole genome phylogeny (Figure 1). Genes are considered present (black) with greater than 80% average translated nucleotide identity, calculated using BLASTx (BLAST+ v2.2.26 (66)), across the total reference gene length. Figure was prepared using SeqFindr (77).

The Locus of Enterocyte Effacement (LEE) was present in all strains within ST106A, ST106B, ST118 and ST182, but absent from all other phylogroup B1 lineages in this study. The core LEE regions (encoding the T3SS) from the 44 strains examined in this study shared greater than 95% nucleotide sequence conservation with the corresponding LEE region from *E. coli* 11368 (O26:H11; Figure 3). The site of integration of the LEE was determined by sequence comparison to known insertion sites, namely *pheU*, *pheV* and *selC* as previously observed in *E. coli* strains 11368 (O26:H11), 11128 (O111:H-) and Sakai (O157:H7), respectively (27). Among the subset of strains for which the LEE insertion site could be determined; n37 and n48 (ST118) contained the LEE carrying the Intimin epsilon variant inserted into *pheV*; strains n10 and n11 (ST106B) contained the LEE carrying the Intimin theta variant inserted into *pheV*; and strains n1, n3, n4, n5, n6, n12, n13, n38, n39, n40 and n42 (ST106A) and n43 (ST182) contained the LEE inserted into *pheU*. The latter two cases likely represent independent events given the evolutionary distance between these two lineages and the differences in the Intimin type (ST106A: Intimin beta; ST182: Intimin epsilon) (Figure 1). The LEE insertion site could not be unequivocally determined in strains n8, n9, n17, n36, n37 and n44 from the draft sequence data alone, although the likely insertion sites could be predicted given the position of the LEE within other strains from the same lineage (Figure 1). Taken together, the data indicate that Stx and/or LEE acquisition has occurred multiple independent times among strains in the B1 phylogroup to give rise to different STEC lineages.

**Figure 3.**
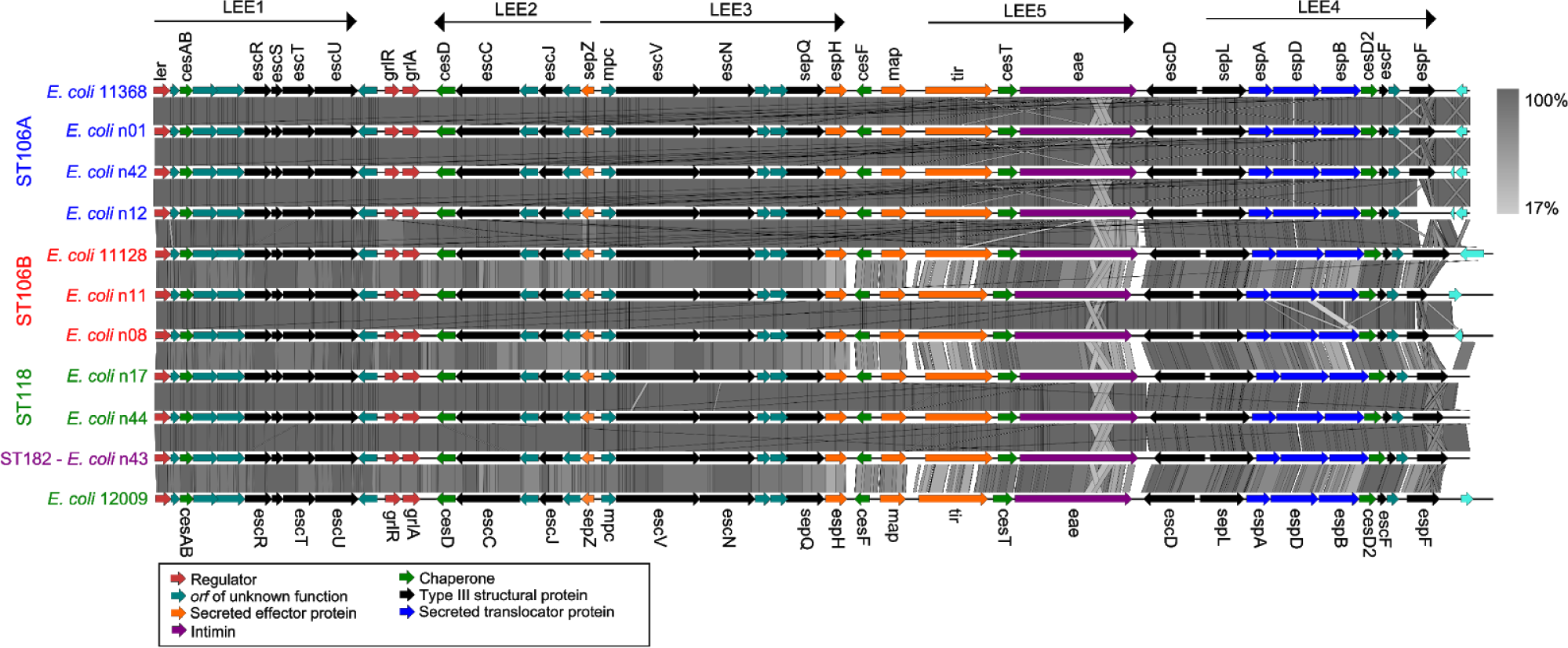
Sequence comparison of LEE region from non-O157 STEC. Translated nucleotide comparison (BLAST 2.2.27+ tBLASTx) (66) of part of the Locus of Enterocyte Effacement (between *ler* and *espF*) in representative non-O157 STEC from different serogroups and sequence types. Strain labels have been color coded according to lineages defined in Figure 1. tBLASTx alignment identity score is indicated by scale gradient. Figure was prepared using EasyFig (71). CDS were color coded according to function outlined in (74).

### Type III secreted effectors

All non-O157 LEE+ STEC were found to encode the six known LEE encoded effector genes (*espG, espZ, espH, map, tir, espF*). In addition, 21 different phage-associated exchangeable effector loci (EELs) were found within the genomes of strains that encoded the LEE, with some variation in effector repertoire between and within lineages (Additional File 2). A number of EELs were consistent with previously defined effector loci (26) and these designations have been included in Additional File 2 to maintain a standard nomenclature. Notably, O121 strain n43 encoded an effector loci with a gene order not previously observed in (26), encoding EspO, EspN, EspM. The largest number of effectors (>30 per genome) were found to be encoded in the ST106 and ST118 lineages, with approximately half this number identified in the O121 strain n43 (Additional File 2). Each EEL (EEL1-EEL21) differs in effector gene content, order and number of effectors. Six of the ten STEC O26:H11 strains within the ST106A lineage are missing one or two EELs when compared to the reference O26:H11 str. 11368 genome. Likewise, the O111:H11 strains within lineage ST106A shared a similar EEL profile as the O26:H11 strains (Additional File 2). In contrast, the O111 strains from lineage ST106B contained several different EELs compared with the ST106A strains (Additional File 2) consistent with the acquisition of different prophages after their acquisition of LEE. Interestingly, some EELs (EEL01, 02, 05, 10) were shared by both ST106A and ST106B lineages, albeit with some minor differences indicative of lineage-specific loss of individual genes in the case of EEL05 (Additional File 2). Examination of other LEE+ STEC strains in our collection identified other EELs shared between phylogenetically distributed strains, including the O121 strain n43 from the ST182 lineage (indicating independent acquisition of the same EEL).

### Other STEC virulence factors

A number of other virulence factors have been associated with STEC pathogenesis in previous studies. We queried the strains examined in this study as well as selected strains from the B1 phylogroup using BLAST and read-mapping for a range of STEC virulence factors, including genes encoding adhesins, autotransporter proteins, fimbriae, cytotoxins and genes from plasmids origin defined pathogenicity islands including the Yersinia high pathogenicity island [HPI], O-island 112 and O-island 43/48 (Figure 2; Supplementary Table 3)

The *iha* and *ehaA* genes have been previously described as well conserved in STEC (28–30). Within this study, all STEC strains were positive for *ehaA*, however *ehaA* was also present in non-STEC strains IAI1, 59899 and E24377A. The *iha* gene was conserved in all strains except for ST234 strains and strains n39, n38 and n51 (ST106A). The distribution of other virulence factors varied, even among strains within the same serotype or sequence type. The O-islands 122 and 43/48 were conserved among LEE positive strains, however *pagC* from the O-island 122 was absent in ST106A and ST118 strains. The *fyuA* gene, used as a marker for the *Yersinia* HPI, was conserved in LEE positive ST106A strains and ST379 strains, but not in ST106B strains or strain n43 (O121).

Several virulence factors have been associated exclusively with LEE negative strains; *saa* (31), *sab* (16), *epeA* (32) and *subAB* (33). The *saa* gene is often associated with LEE negative strains that originate from ruminants, but has not been associated with clinical STEC identified from HC or HUS patients. The *saa* gene was absent from all strains examined in this study. The *sab*, *epeA* and *subAB* genes were originally identified on a large plasmid carried by an STEC O113 strain; these genes are conserved in all O113 strains (ST234). These genes were also identified in strain n47 (ST650), suggesting that n47 carries a similar plasmid.

Genes associated with the pSAK virulence plasmid, namely *espP*, *katP*, and *hlyA*, were present in ST106A and ST106B strains with some exceptions. For example, the pSAK associated genes, *espP, katP*, and *hlyA*, are present in n02 (ST461 O91:10) and n48 (ST118 O103:H2) but absent from other strains of the same sequence type. Conversely, all strains within ST106B possessed the same plasmid-associated genes except n10 (O111:H8). This plasmid profile is consistent with an ancestral acquisition of a pSAK-like plasmid prior to divergence of ST106A and ST106B lineages, with subsequent sporadic strain-specific loss. However, we cannot rule out multiple independent acquisition without complete sequencing of the plasmid content of these strains. All strains were negative for the *aggA*, *aggC* and *aggR* genes identified in the plasmid carried by 2011 O104:H4 outbreak strains.

### Comparison of serogroup and phylogenomic approaches

O-antigen serotype was lineage specific in ST106B (O111), ST379 (O128) and ST234 (O113) strains. In contrast, O-antigen serotype was inconsistent with whole genome phylogenomics and sequence typing in three instances involving O91, O103/O45,and O111/O26 strains, indicating widespread lateral gene transfer (LGT) of the O-antigen biosynthesis genes. O91 strains clustered into three separate lineages in the whole genome phylogenetic tree (Figure 1): (i) O91:H10 strains belonged to the ST461 lineage; (ii) O91:H21 strains belonged to the ST89 lineage which also contained a distinct O-antigen untypeable H21 strain (n41); (iii) the O91:NM strain n32 was typed as ST815 and did not cluster with any other STEC strains.

While O45 and O103 have been described as distinct members of the ‘top-six’ non-O157 serogroups by the Centre of Disease Control and Prevention (34), this work classifies these two serogroups within a single lineage (ST118) (Figure 1). The ST106A sub-lineage included O111:H11 and O26:H11 strains, and was distinct from the O111:H8,O111:NM, and O111:H-strains that comprised the ST106B sub-lineage. This indicates that O111:H11 and O26:H11 strains share a common ancestry, and that the O26 O-antigen region was most likely acquired by lateral gene transfer in ST106A.

Serotype was determined by nucleotide alignment of the region between the *gnd* and *galF* genes in the O antigen biosynthesis locus. Regions with the same O-antigen serotype possessed >98% sequence identity over the O-antigen biosynthesis region, whereas regions with different O-antigen sequences shared no significant nucleotide conservation and very low (7-26%) amino acid identity. This was also observed between the most closely related strains that showed evidence of O-antigen lateral gene transfer, namely O26 and O111 strains from ST106A and ST106B (Supplementary Figure 3).

O-antigen typing was also validated by sequence comparison of the *gnd* gene, which encodes 6-phosphogluconate dehydrogenase, and is located immediately upstream of the O-antigen locus. This approach has previously been used for molecular-based serogrouping of STEC (35), where *gnd* sequences with >99% nucleotide conservation define the same O-antigen type (36). Indeed, a Maximum-likelihood consensus tree based on the *gnd* gene sequence (Supplementary Figure 4) corresponded with O-antigen type (Additional file 1) for most of the STEC strains examined. The exception was n32 (O91:NM), which did not cluster with other O91 strains or any other STEC strain. BLAST comparisons showed that the *gnd* sequence from n32 shared, on average, 96.38% nucleotide conservation between other O91 strains and 95.52% nucleotide conservation with other STEC. Comparison of the sequence of the entire O-antigen region in n32 to other O91 and STEC showed the same level of conservation (Supplementary Figure 5). These data suggest that n32 may have acquired the genes encoding the O91 O-antigen region through lateral gene transfer independent of the *gnd* gene. However, this would suggest that *gnd* typing may not be suitable for STEC as described previously (35), which would hamper the utility of using *gnd* typing as a proxy for O-antigen typing.

**Figure 4.**
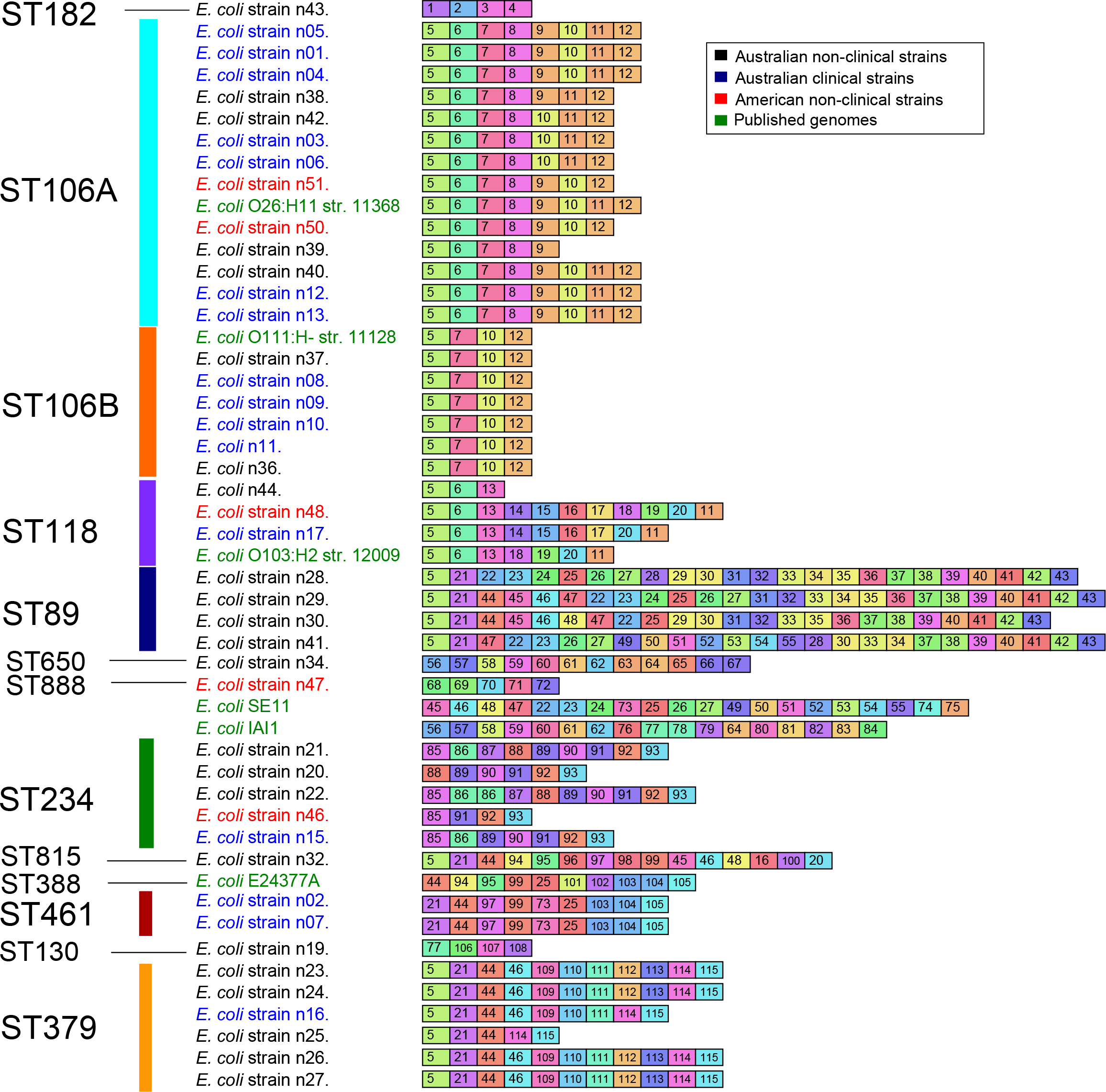
Conservation of CRISPR1 spacers in non-O157 STEC and B1 phylogroup *E. coli*. Graphic representation of spacer content from CRISPR1 for B1 phylogroup *E. coli* including strains from this study and other B1 phylogroup *E. coli* that share spacer sequences including SE11, E24377A and IAI1. A uniquely colored box and symbol combination designates each spacer sequence. Sequences are listed (left to right) from farthest to nearest the CRISPR leader sequence. Strains and lineages are listed in order and colored according to the scheme used for the phylogram in Figure 1.

### CRISPR diversity within non-O157 STEC

The diversity of CRISPR arrays within non-O157 STEC strains was explored to determine the suitability of CRISPRs as a typing method (37). *E. coli* and *Salmonella* can have two CRISPR loci, with the CRISPR associated (*cas*) genes at each locus variable and classified as either I-F or I-E *cas* subtypes (38,39). Up to two CRISPR arrays flank each *cas* loci at a specific insertion: for *E. coli* CRISPR/*cas* these are designated CRISPR1 (between *cysD-cysH*) and CRISPR2 (between *cysH-ygcF*); for Ypest CRISPR/*cas* subtypes these are designated CRISPR3 and CRISPR4 (between *clpS-aat*).

The conservation of spacer sequences was examined in the 44 STEC strains as well as the representative *E. coli* strains. CRISPR spacers from the B1 phylogroup STEC strain spacer repertoire were also found in *E. coli* strains E24377A, B REL 606 and IAI1, but were not found in other *E. coli* (data not shown). In turn, spacer sequences from other *E. coli* did not appear in B1 phylogroup STEC strains.

Each locus varied in gene content and spacer sequences for all STEC strains, even among strains within the same lineage. All 44 STEC strains contained a CRISPR1 locus, with a total of 115 unique spacers identified across the B1 phylogroup. The majority of these were localized in clusters of 3 to 25 spacers, with a median of 8 spacers per CRISPR1 locus (Figure 4). There was little similarity in spacer content between STEC strains from different lineages. For example, no spacer sequences from LEE-positive strains were found in LEE-negative strains. Some variation of spacers within a particular lineage was observed and could be due to strain or lineage specific deletion of spacers. Some spacer sequences were consistent across strains of the same lineage and may represent potential genotyping targets to identify individual STEC lineages.

CRISPR2 arrays were detected in the majority of strains in this study, localized blocks comprising multiple clusters of 2 to 4 spacers, with a median value of 9 (Supplementary Figure 6). The diversity of spacer sequences was variable within different lineages; strains n05, n01, n04, n38 (ST106A) contained common spacers but were distinct from other ST106A strains n03, n06, n50, n39, n40, n12 and n13. Strain n51 possessed a set of 9 spacer sequences that were not present in any other ST106A strains. Similarly, ST106B was separated into two groups, with strains n37 and n08 containing one set of spacers and strains n9, n10, n11 and n36 containing a different (but conserved) set of three spacers. Spacers were conserved within ST118, except for strain n17, which carried two spacers that were not detected in any other B1 phylogroup strain examined in this study. Some lineages showed a high degree of conservation of CRISPR2 spacer sequences with examples of step-wise spacer acquisition (ST234) or deletion (ST379). Further resolution of individual lineages will assist in determining the significance of spacer diversity within the STEC population.

In terms of CRISPR3 and CRISPR4, CRISPR3 was only detected in the n34 genome, with two arrays of 13 and 22 spacer sequences identified within the region between *clpA* and *infA*. Strain n34 also carried a full set of Ypest *cas* genes. Instead of the CRISPR3 array, strains n1, n4, n38, n39, n40, n12 and n13 carried a mobile genetic element encoding an integrase, a number of genes encoding hypothetical proteins and genes encoding a YeeU/YeeB toxin/anti-toxin system. No CRISPR array was detected in the CRISPR3 loci for any other B1 phylogroup STEC strains examined in this study. Instead, these strains contained genes encoding a tRNA-Ser and a Translation initiation factor IF-1 at this locus. All non-O157 strains examined in this study lacked CRISPR4 and no novel CRISPR regions were found using whole genome detection of CRISPR arrays with either PILER-CR or CRISPRFinder.

### Genomic features shared by STEC strains

Whole genome alignment was utilized to identify genomic regions conserved among STEC genomes. We identified 22 genomic regions that were predominantly associated with STEC strains, eight of which were determined from the whole genome alignment as conserved across all STEC strains used in this dataset. These regions are summarized in Table 2, together with details of their location relative to the published 11368 (O26:H11) and EDL933 (O157:H7) genomes. Of the 22 regions identified by this analysis, three large regions (>5kb) were present across all twelve representative STEC strains, but not present in K12 MG1665 (Table 2, bold): (i) a 10,117bp region encoding the Type VI Secretion system as part of O-island 7; (ii) genes encoding CRISPR associated genes associated with the CRISPR1 region, and (iii) the second cryptic T3SS (ETT2) within O-island 115.

**Table 1.**
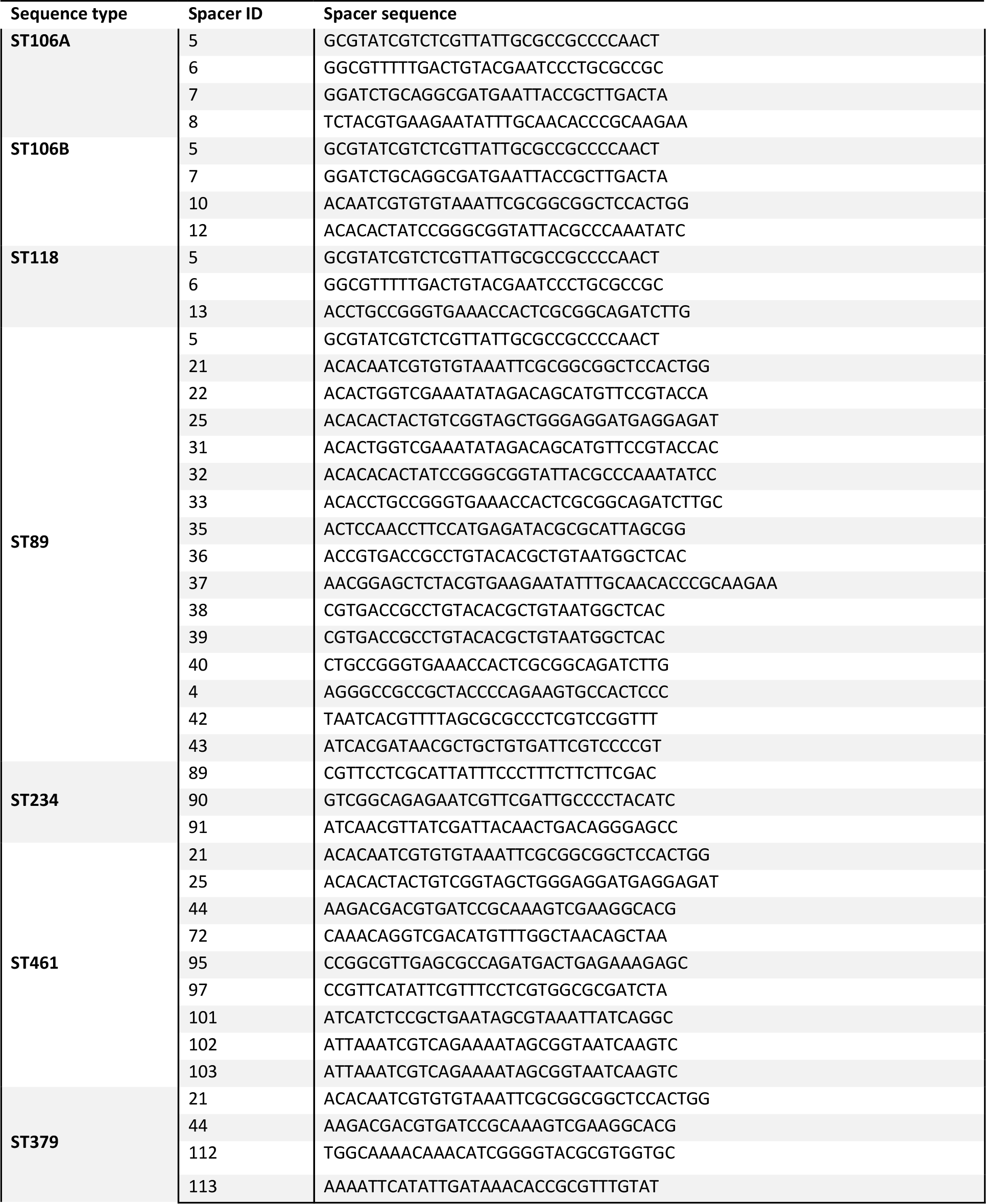
CRISPR spacer sequences conserved within each lineage

**Table 2.** Genomic region conserved across STEC

| Conserved<sup>1</sup> | O26 Start | O 26 Stop | O26 Length | O26 Locus Tags | O157 EDL933 Locus Tags | Description<sup>2</sup> |
| --- | --- | --- | --- | --- | --- | --- |
| * | 21257 | 22142 | 885 | ECO26_0022 | Z0025 | Partial match to "T3SS effector-like protein EspX-homolog". (O-island 1) |
| * | **248034** | **258211** | **10177** | **ECO26_0220-ECO26_0230** | **Z0250-Z0258** | **Macrophage toxin and Type VI secretion system (O-island 7)** |
|  | 312255 | 312976 | 721 | ECO26_0291-ECO26_0292 | Absent | 'Conserved predicted protein' between *yafP* and *pepD*. Absent in EDL933. |
|  | 640740 | 645802 | 5062 | ECO26_0611-ECO26_0619 | Z1888-Z1896 | Structural phage related proteins (O-island 52) |
|  | 658907 | 659790 | 883 | ECO26_0630 | Z1930 | Putative hydrolase, phage related (O-island 52) |
|  | 669640 | 670733 | 1093 | ECO26_0636 | Z0700 | Putative receptor and insertion sequence:IS677 (O-island 30) |
|  | 675294 | 677991 | 2697 | ECO26_0639-ECO26_0641 | Z0705-Z0707 | Unknown (Rhs related) and VgrG protein (O-island 30) |
| * | 695271 | 695387 | 116 | ECO26_0656 | Z0722 | HokC – small toxic membrane peptide |
| * | 1052404 | 1054915 | 2511 | ECO26_1001-ECO26_1003 | Z1108-Z1110 | Conserved putative proteins and aquaporin AqpZ, (O-island 42) |
| * | 1624945 | 1625425 | 480 | ECO26_1653 | Z1844 | Hypothetical protein (Phage related) Cryptic prophage CP-933C |
|  | 1740868 | 1741751 | 883 | ECO26_1769-ECO26_1771 | Z2036-Z2037 | Integrase, excisionase and exonuclease, Phage related (O-island 57) |
|  | 1746722 | 1747229 | 507 | ECO26_1779-ECO26_1780 | Z2046-Z2047 | Phage repressor and anti-repressor (O-island 57) |
| * | 2003329 | 2003974 | 645 | ECO26_2057 | Z2262 | VgrE gene – Rhs related (O-island 65) |
|  | 2052881 | 2053623 | 742 | ECO26_2092-ECO26_2093 | Z2213-Z2214 | Predicted peptidase and porin protein |
|  | 2487136 | 2487833 | 697 | ECO26_2563 | Z3664 | putative IS609 transposase TnpB (O-island 103) |
|  | 2488392 | 2488884 | 492 | ECO26_2564 | Z3665 | putative IS609 transposase TnpA (O-island 103) |
|  | 3445476 | 3446301 | 825 | ECO26_3520-ECO26_3521 | Absent | Between *ypfJ* and *purC* : predicted post-segregational-killing toxin/anti-toxin |
| * | 3746864 | 3747995 | 1131 | ECO26_3806-ECO26_3807 | Z4045-Z4046 | putative 4-hydroxybenzoate decarboxylase (O-island 110) |
| * | **3761305** | **3770325** | **9020** | **ECO26_3823-ECO26_3832** | **Z4062-Z4071** | **CRISPR 2 – Cas genes** |
|  | **3885013** | **3892930** | **7917** | **ECO26_3931-ECO26_3944** | **Z4180-Z4190** | ***E. coli* Type Three Secretion System 2 (ETT2) Cryptic Type III secretion system (O-island 115)** |
|  | 4167893 | 4173254 | 5361 | ECO26_4197-ECO26_4200 | Absent | Predicted pillin, usher and fimbrial protein |
|  | 5024670 | 5026291 | 1621 | ECO26_4995 | Z5029 | Putative adhesin (O-island 144) |

<sup>1</sup>Conserved across all STEC sequenced in this study
<sup>2</sup>Bold rows: Regions not present in Lab-adapted K12 MG1665

In contrast to the diversity observed within *E. coli* CRISPR spacer sequences, the CRISPR associated genes (*cas*) within CRISPR1-2 were conserved across all but one STEC strain (STEC_7v) examined in this study, as well as previously published STEC complete and draft genomes from a range of *E. coli* phylogroups (E, B1, B2, D) (Additional File 3). Cas genes were *cas* subtype I-E based on gene order and chromosomal location, but closer inspection with sequence comparisons showed variation within *cas* genes. When considering representative genomes across the *E. coli* species (Additional File 3), the CRISPR1 *cas* loci could be classified into three variant alleles according to its similarity with representative regions from CFT073 (from which it is absent), MG1655 (K12), or 11368 (O26:H11) (Additional File 3). The *cas* genes in MG1655 and 11368 were divergent at the amino acid level, with 14% amino acid identity on average between orthologous proteins encoded by *casA-E* (Supplementary Figure 7). The STEC O26:H11 *cas* variant was found in all strains within B1 phylogroup, including B1 phylogroup STEC and other *E. coli* such as ETEC E24377A, EAEC strain 55989 and commensal strains SE11 and IAI1. The STEC O26:H11 *cas* variant was also conserved in O157:H7 strains and n43, the ST182/O121 STEC strain that belongs to a divergent STEC lineage.

## Discussion

Non-O157 STEC are increasingly recognized as an important food-borne pathogen responsible for global outbreaks (40). While O157:H7 remains the dominant serotype in the United States, United Kingdom and Japan; STEC serogroups including O26, O111, O103, O121, O45,O128, O91 and O113 have also been associated with human disease (8,41). Strains from these serogroups include both LEE positive and LEE negative variants (14,42). This highlights the need for wider studies into STEC other than the dominant O157:H7 serotype. To address this, we have analyzed the whole genome sequence of 44 non-O157 STEC strains from different origins to explore their overall diversity.

We aimed to examine the whole genome phylogeny of the most prevalent non-O157 STEC serogroups including LEE positive and LEE negative strains. It has been previously noted that STEC is comprised of multiple lineages that have evolved from parallel paths (25,26,43). The phylogenomic analyses presented here placed all strains within the B1 phylogroup (53) and confirmed the previously reported distinction between LEE positive O26, O111, and O103 strains (26). We also demonstrate that LEE negative serogroups of clinical importance, namely O91, O113 and O128, form separate lineages within the B1 phylogroup. Our analysis of the LEE positive O121 strain showed that it was distinct from all other STEC strains, consistent with previous studies of O121 (45).

Defined STEC lineages were largely concordant with EcMLST typing (46) and we referred to lineages in accordance to that scheme to allow for a scalable nomenclature. ST106 correspond to the previously defined EHEC-2 clonal group, ST118 corresponds to the STEC-2 clonal group and ST182 correspond to the distant STEC-3 clonal group (53,54). However, ST106 strains comprised two individual sub-lineages (Figure 1) and were designated as ST106A, which comprised H11 strains of both O26 and O111, and ST106B, which included O111 strains of flagella types other than H11. A number of singleton STEC lineages were also identified in this study with many more lineages expected as the genomes of further non-O157 STEC are sequenced.

Serotyping is a standard classification method for *E. coli*, particular for STEC/EHEC. Typing schemes such as seropathotyping (10) have linked disease potential of EHEC strains to particular serotypes based on epidemiological and prevalence studies. Previous studies have shown that certain serogroups (such as O174) are distributed across different phylogenetic backgrounds, consistent with lateral gene transfer of the O-antigen region (47). Within our study of 44 STEC strains, we observed several instances where O-antigen was inconsistent with the defined phylogeny, suggesting that lateral transfer of the O-antigen region may be more prevalent than first suspected. In all cases, O-antigen typing was verified through sequence comparison of the O-antigen regions. Most notably, we could distinguish O111 strains based on flagella type that placed O111:H11 strains (n12 and n13) within the ST106A lineage, separate to other O111 strains in ST106B. The O111 antigen locus has already been associated with lateral transfer between O35 in *Salmonella* (48), and the mobility of this region would explain conflicting virulence profile results within O111 strains. Other O26:H11 and O111:H11 strains share common ancestry, based on genomic content (49). O103:H11 strains were also found to be more closely related to O26:H11 rather than other O103 strains (50), highlighting the need to examine the phylogenetic background of O111, O103 and O26 strains when comparing virulence factor profiles.

We also found that O91 strains in our study belonged to separate lineages within the determined phylogeny. A previous MLST-based study has shown that O91 strains of different sequence types had differential associations with HC or HUS (51). Our phylogenomic analysis suggests that the acquisition of genes encoding O91 antigen has occurred through multiple independent events, perhaps driven by host adaptation rather than acquisition by a common ancestor. Similarly, an O45 serogroup strain was found to be almost indistinguishable at the core genome level to three O103 strains, including the reference strain 12009 (26). These results highlight the importance of a sequence-based genotyping approach combining lineage and virulence gene content for the routine identification of STEC strains.

The T3SS encoded on the LEE is essential for the production of attaching and effacing lesions by EHEC and EPEC, and its presence is associated with clinically dominant STEC strains (10). The acquisition of the LEE in EHEC and EPEC is considered a key evolutionary event for both pathotypes (25). LEE positive STEC strains with in this study were distributed across four lineages with differential insertion sites and Intimin subtypes (ST106A: Beta-*pheU*, ST106B: Theta-*pheV*, ST118: Epsilon-*pheV* and ST182: Epsilon-*pheU*). LEE positive STEC O26 was suggested to have arisen through step-wise evolution from atypical EPEC O26 strains, whereby the LEE was inserted into *pheU*, and the *stx1* or *stx2* genes were gained through phage integration on the chromosome (52). In contrast, O111 strains are thought to have independently acquired the LEE in *pheV* (53,54). Our data suggest that the LEE has been acquired within STEC on at least five individual occasions (O157, ST106A, ST106B, ST118 and ST182). The effector sequence profile of complete genomes from LEE positive strains suggest that LEE acquisition was accompanied by phage-mediated lateral gene transfer of a distinct, lineage-specific effector repertoire (26,55). Phage-associated effectors were not identified in any LEE negative strains, suggesting that LEE is necessary for selection and maintenance of EELs. Intriguingly, we were able to discern several EELs in common between the different lineages, with different locations suggesting independent acquisition of common genetic elements. Furthermore, the observation of EELs with common locations between ST106A and ST106B which have separate LEE insertion sites, suggest that a LEE may have been acquired prior to divergence of these lineages, followed by displacement with an independently acquired LEE pathogenicity island in one of these lineages after divergence. The sequencing of multiple STEC genomes using long-read technologies should enable a detailed analysis of LGT events that have occurred along each STEC lineage.

Stx type was heterogeneous within ST89, ST234 and ST118 lineages. It is possible that the *stx* genes had been lost during cell culture or infection as observed in previous studies (56,57). Stx2d_act_ (elastase-cleaved), was only found in LEE negative STEC strains within the ST89 and ST379 lineages. In other studies, Stx2d_act_ has been found to be prevalent within LEE negative strains (58,59).

CRISPR have been identified in a number of bacterial species (38,60) and their distribution has been linked with phylogenetic grouping in *E. coli* (61). CRISPR have been shown to have a distinct role in phage defense in *E. coli* (58), and the modification of CRISPRS/*cas* genes can alter susceptibility to infection in some species (59). Previously, CRISPR loci were proposed as a method to distinguish between highly virulent STEC serotypes (O157:H7, O26:H11, O145:H28, O103:H2, O111:H8, O121:H19, and O45:H2), as the spacer sequences within these loci are unique to these STEC (37). However, *E. coli* sero-pathotypes in general have shown little association to CRISPR content (60,61). This was also reflected in our study, which showed that spacer repertoire does not distinguish EHEC from non-EHEC strains, suggesting that any CRISPR based scheme to define EHEC from other *E. coli* would not be feasible without an additional method such as *stx-*typing. However, we did find that spacers within CRISPR1 could differentiate individual STEC clonal lineages. Therefore, it may be possible to develop a scheme to differentiate individual STEC clonal lineages using CRISPR1 loci.

Comparisons within *E. coli* have yielded little correlation between CRISPR arrays and resistance to foreign elements (62), but this does not incorporate the sequence variations noted here within the *cas* complex. Our results present the intriguing possibility that the types of *cas* genes carried by STEC are directly related to the observation that only particular *E. coli* lineages are able to acquire phage that carry the *stx* gene or genes encoding Type III effectors.

## Conclusions

In conclusion, while pathogenic bacteria are usually discussed in the context of lineage specific virulence and other gene acquisition, here we present STEC as a set of diverse lineages evolving in parallel with independent acquisition of virulence factors. Using the phylogenomic analysis presented here, we have compared alternative typing methods such as serotyping, EcMLST and CRISPR typing, and found that each of these approaches have their own shortcomings in representing the divergent nature of STEC. Given, the varied distribution of STEC virulence, we also attempted to define the CRISPR-cas locus as a common genomic element that could provide the genomic background for Stx-encoding phage acquisition. While this work presents many broad findings, additional insights a more in-depth study of variation within and between the complete genome sequences of a more geographically diverse representative STEC isolates.

## Supporting information

Additional File 1 - Strains in this study

Additional File 2 - FullEELTable

Additional File 3 - Distribution of CRISPR1 cas genes

Addtional File 4 - Effector profiles

## Methods

### Bacterial strains

Forty-four STEC strains from collections held by Queensland Health Forensic and Scientific Services (QHFSS) (n=16), The Commonwealth Scientific and Industrial Research Organisation (CSIRO) (n=23) and Washington State University (WSU) (n=5) were used in this study. These strains encompass the major non-O157 serogroups and include strains from human, ruminant livestock and contaminated food sources (Additional File 1).

### Genome sequencing and annotation

All strains were sequenced on the Illumina HiSeq2000 platform. The resulting paired end 100 base pair reads (average of 302 bp insert size with standard deviation of 108 bp) were filtered using PRINSEQ-lite. Reads were trimmed at both ends to achieve a mean quality cut-off of Q20 and minimum read length of 80 base pairs. Filtered reads were assembled using SPAdes version 2.5.0 (63) with default kmers (21, 33 and 55) and with inbuilt read and scaffold correction, and the “--careful” flag. The resulting assemblies included a subset of low coverage scaffolds, which were artifacts of the sequencing and assembly process. These low coverage scaffolds could be partitioned from scaffolds with an expected coverage, and were filtered out using a coverage cut-off calculated independently for each assembly. The cut-off was based on scaffolds average coverage while adjusting for GC bias and had an average of 10 with a standard deviation of 5. In contrast, the average read coverage for each genome was 252 with a standard deviation of 134. The filtered, assembled scaffolds were ordered using Mauve ContigMover (mauve_snapshot_2012-06-07) (64) against the published *E. coli* O111:H-strain, 11128 (Accession no. AP010960), and annotated using Prokka version 1.5.2 (65). Genome data are available on EnteroBase at http://bit.ly/AlikhanetalSTEC

### Phylogenetic analysis and recombination testing

To compile a set of core gene sequences for subsequent phylogenetic analysis, we first retrieved all predicted gene sequences from the published complete genome of *E. coli* O111:H-str. 11128 (Accession No. AP010960), and identified homologs in available complete *E. coli* genomes from the B1 Phylogroup, non-O157 STEC and O157:H7 Sakai using the Basic Local Alignment Search Tool (BLAST) (version 2.2.26+) (66). Putative homologs for each reference gene were defined as the predicted gene with the best scoring BLAST alignment match that had greater than 90% nucleotide identity over 90% gene length to the respective reference gene from strain 11128. Each cluster of homologous genes was subsequently aligned using MUSCLE (67) and then concatenated into a single alignment sequence. Variable sites were extracted from this alignment to produce a concatenated and aligned sequence of SNPs for each of the taxa in PHYLIP format for phylogenetic analysis. This approach was implemented through an in-house script Dryad (68). The aligned SNP sequence was used in PhyML (v20120412) (69) to infer a maximum-likelihood phylogram using the HKY85 substitution model and 400 bootstraps.

To assess if recombination within core gene families impacted on the topology of the tree, a strict filter was imposed such that ortholog groups were removed if they exhibited significant evidence for recombination (p <0.05) for at least two out of three homoplasy tests (NSS, MaxX^2^ and PHI) implemented in PhiPack (70). Variable sites were extracted, and a phylogenetic tree was determined using the same methodology as described above.

### Sequence type and serogroup determination

Sequence types were defined in accordance with the *E. coli* MLST (EcMLST) scheme (46). Gene sequences were aligned using nucleotide-to-nucleotide BLAST (BLASTn v2.2.26+) (66), and exact matches were used to identify each allele variant. Allele profiles were queried against the EcMLST database to determine sequence type. The serogroup of each strain was confirmed *in silico* by comparing the nucleotide sequences, using BLAST (BLASTn v2.2.26+), of a region within the LPS biosynthesis locus (between *hisG* and *yegH*) for O-antigen typing and the *fliC* gene for flagella typing. Comparisons of the O-antigen biosynthesis region were visualized using EasyFig (71).

### Stx, LEE and effector profiling

Shiga toxin encoding genes were detected through amino acid and nucleotide sequence comparison (BLASTp and BLASTn) to Stx2a (protein id: CAA71748) and Stx1a (protein ID: AAG57228) sequences from *E. coli* O157:H7 str. EDL993. Detected *stx* subunit B gene sequences were compared with BLASTn to the GenBank nucleotide non-redundant database and required an identical nucleotide sequence match to previously sequenced *stx* genes from STEC strains. *stx* subtypes can vary by as little as a single SNP and isolates harboring multiple *stx2* genes can have these genes interpreted as a collapsed repeat in assembled contigs. To address this, *stx* copy number and type was determined through the analysis of SNPs from the *stx* genes generated from mapping reads onto the O157:H7 Sakai genome (Accession no. BA000007) with BWA (72). *stx* copy number and type was also confirmed with Mapsembler (73). The subtype for matching strains was defined in accordance with existing literature (Supplementary Table 3).

The LEE insertion site for sequenced strains was determined through genome comparison to known LEE insertion sites as defined in (74), using the Artemis Comparison Tool (75); *yicK* to *selC*, observed in O157:H7 Sakai (Accession no. BA000007), *yghD* to *pheV*, observed in O111:H-11128 (Accession no. AP010960) and O103:H2 12009 (Accession no. AP010958), and *cadC* to *pheU* observed in O26:H11 11368 (Accession no. AP010953).

Effector repertoires were annotated in each draft genome using the EffectorFAM database of profile HMMs built from confirmed effector families, including all *E. coli* Sakai O157:H7 effectors described by Tobe et al. 2006 (55) (see Additional file 4) (76). Genomic context of each effector was carried out by ACT comparisons with representative complete genomes from each LEE+ lineage (i.e. O26:H11 str. 11368 for ST106A, O111:H-str. 1128 for ST106B, and O103:H2 str. 12009 for ST118).

### Virulence profile

Virulence factor profiles across *E. coli* genomes were generated using SeqFindr (77). Virulence factors were considered present with greater than 80% average nucleotide identity across the total reference gene length. Comparisons between individual genomes and verification of SeqFindr results were performed using BLAST+ (v2.2.26+) (66), Artemis Comparison Tool (75) and EasyFig (71).

### CRISPR detection

Genomes were interrogated for CRISPR spacer sequences using PILER-CR (78) and verified with CRISPRFinder (79) across the whole genome. CRISPR loci defined in (62) were also inspected using nucleotide-to-nucleotide BLAST (BLASTn v2.2.26+)(66), Artemis Comparison Tool (75) and EasyFig (71). The distribution of unique spacer sequences was visualized using binCrisp, a custom python script developed as part of this study (80).

### Whole genome comparison

The sequences of forty-four non-O157 STEC genomes sequenced as part of this study and 11 representative strains from the major *E. coli* pathotypes and phylogroups for which the complete genome was available were aligned using Mugsy version 1.3 (81)(Supplementary Table 2).

Alignment blocks from the whole genome alignment were filtered to identify putative STEC-specific regions. Alignment blocks required no corresponding match in K12 MG1665 or HS but were conserved in twelve STEC genomes chosen as a cross-section of the STEC lineages defined in this study. These included published genomes O26:H11 11368, O157:H7 Sakai, O103:H2 12009 and O111:H-11128 and strains sequenced as part of this study; n10 (O111:H8), n01 (O26:H11), n02 (O91:H10), n28 (O91:H21), n15 (O113:H21), n17 (O103:H21), n16 (O128:H2) and n43 (O121). Accession codes can be found in Supplementary Table 2 and Additional File 1. BLAST (BLASTn v2.2.26+) comparisons were used to verify that regions were conserved across all STEC strains (66).

## Declarations

### Ethics approval and consent to participate

Not applicable

### Consent for publication

Not applicable

### Availability of data and material

Sequenced reads were deposited into the Sequence Read Archive under BioProject PRJEB5529. Accession codes for individual samples can be found in Additional File 1. Genome data are available on EnteroBase at http://bit.ly/AlikhanetalSTEC

### Authors’ contributions

NLB, JAG, SAB performed and compiled effector protein analysis. MSC assisted with virulence profile analyses. All other analyses were performed by NFA. NFA and SAB led the writing of the manuscript, along with NLBZ, NKP and MAS. SAB, RC, TJM, and MAS conceived the study and edited the manuscript.

## Acknowledgements

Authors wish to thank collaborators at Queensland Health Forensic and Scientific Services, The Commonwealth Scientific and Industrial Research Organisation and Washington State University for providing the strains, and Australian Genome Research Facility for sequencing.

