## Supplementary materials and figures for "Multiple evolutionary trajectories for non-O157 Shiga toxigenic *Escherichia coli*"

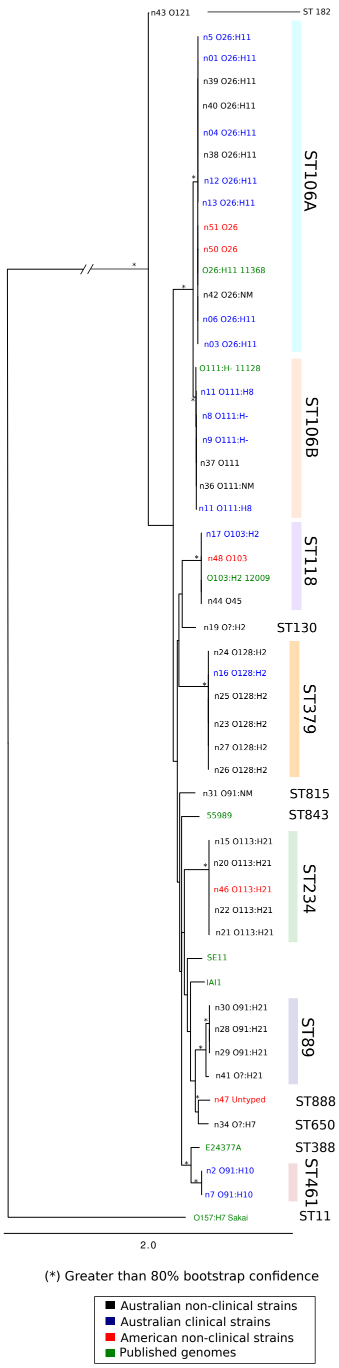

**Supplementary Figure 1.** Phylogenetic relationship of non-O157 STEC and *E. coli* within B1 phylogroup using genes identified as non-recombinant.

Maximum Likelihood (ML) phylogram with asterisks indicating bootstrap support greater than 90% from 400 replicates. The phylogram includes forty-four Shiga toxin positive *E. coli* from this study and seven other *E. coli* from the B1 phylogroup as outlined in the Figure 1 legend. Genomes have been annotated and highlighted according to sequence type defined by the EcMLST seven allele schema, and isolate sources are indicated in the key. Phylogram was constructed using PhyML [3] (HKY85 model) based off 17,438 concatenated SNPs of 1,136 conserved *E. coli* genes that did not show recombination in two out of three tests from PHIpack (Max2, NSS and Phi) [1].

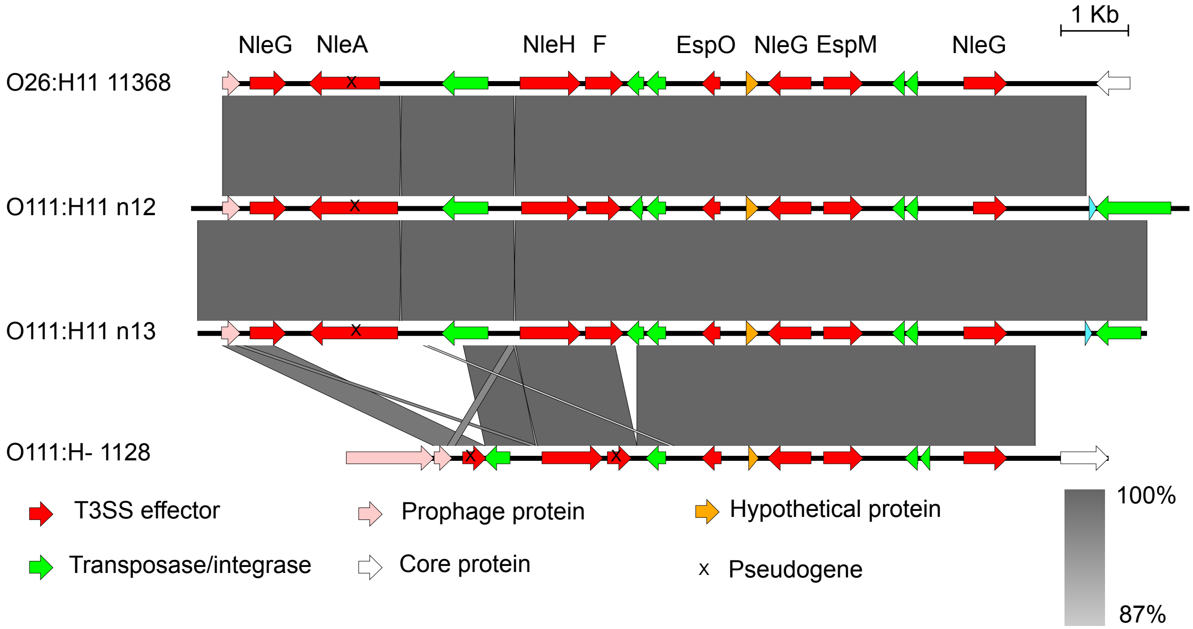

**Supplementary Figure 2.** Organization of effector genes in EEL5.

EEL5 from the two STEC O111:H11 genomes compared to EEL5 from *E. coli* O26:H11 11368 and EEL14 from *E. coli* O111:H- 1128. The grey shading indicates regions of conserved nucleotide sequence determined with BLASTn. The coloured arrows represent genes; pseudogenes are indicated with a slash through the red arrow.

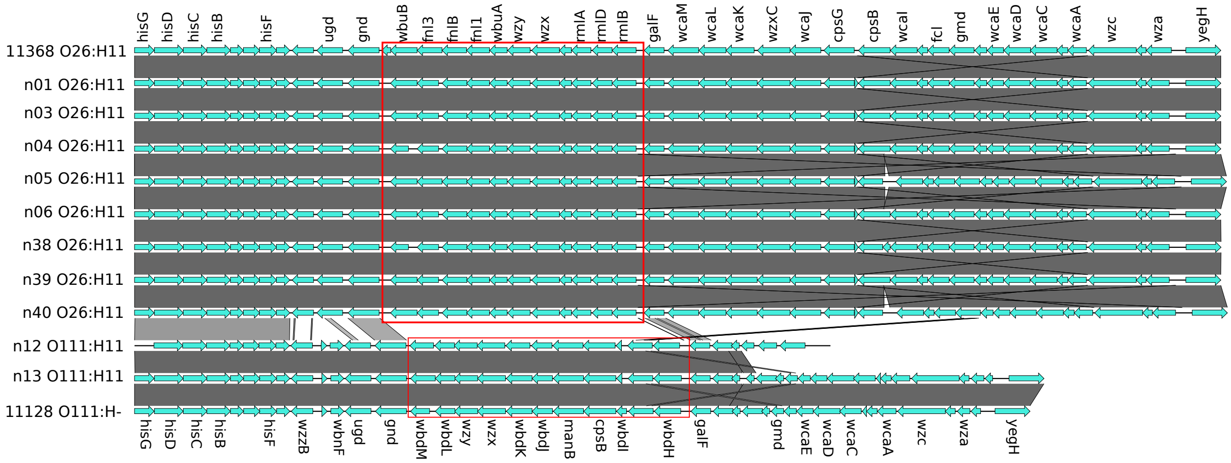

**Supplementary Figure 3.** Comparison of O-antigen synthesis region in ST106A and ST106B strains

Nucleotide to nucleotide BLAST (BLASTn) comparison of O-antigen synthesis region (boxed) between *hisG* and *yegH* from non-O157 STEC strains from ST106A and ST106B. O-antigen region from ST106A strains, n12 and n13, matched O-antigen region from published O111:H- ST106B strain 11128. BLASTn alignment identity score is indicated by scale gradient. Figure was prepared using EasyFig [5].

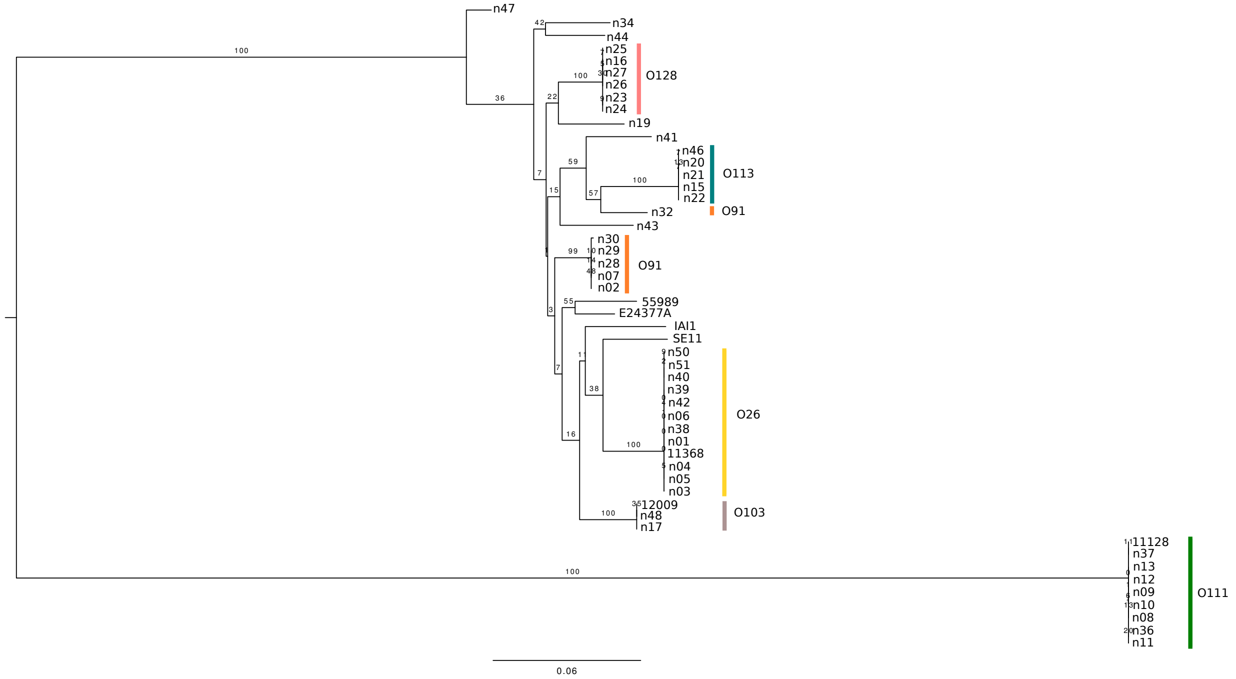

**Supplementary Figure 4.** Phylogenetic relationship of gnd

Maximum Likelihood (ML) phylogram, based on *gnd* gene sequence, from 1000 replicates. The phylogram included forty-four STEC from this study and seven other *E. coli* strains from the B1 phylogroup including EAEC 55989, SE11, IAI1, ETEC E24377A, and previously sequenced non-O157 strains; 11368, 11128 and 12009. Accession numbers of these strains are listed in Table XXX. Genomes highlighted according to O-antigen type. The final figure was prepared with FigTree (v1.4) (http://tree.bio.ed.ac.uk/software/figtree/). The phylogram was built using RaXML [4] with the GTR substitution model.

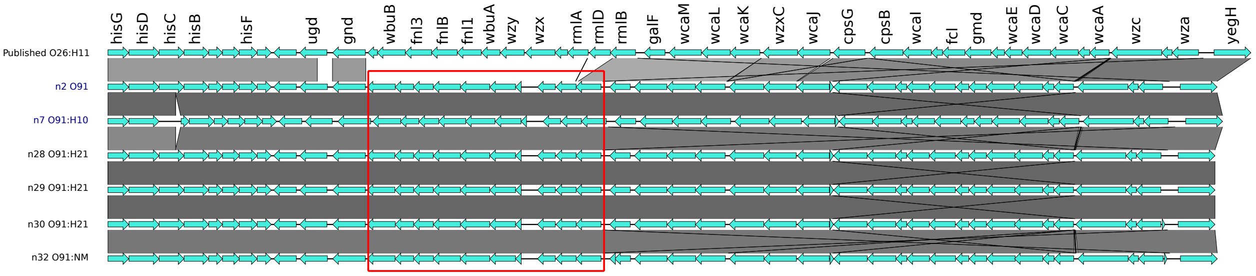

**Supplementary Figure 5.** Comparison of O-antigen synthesis region in O91 strains

Nucleotide to nucleotide BLAST (BLASTn) comparison of O-antigen synthesis region (boxed) between *hisG* and *yegH* from O91 STEC strains. O-antigen regions for O91 strains were identical, while a comparison to O26:H11 strain 11368 (blue) showed no detectable nucleotide similarity. BLASTn alignment identity score is indicated by scale gradient. Figure was prepared using EasyFig [5].

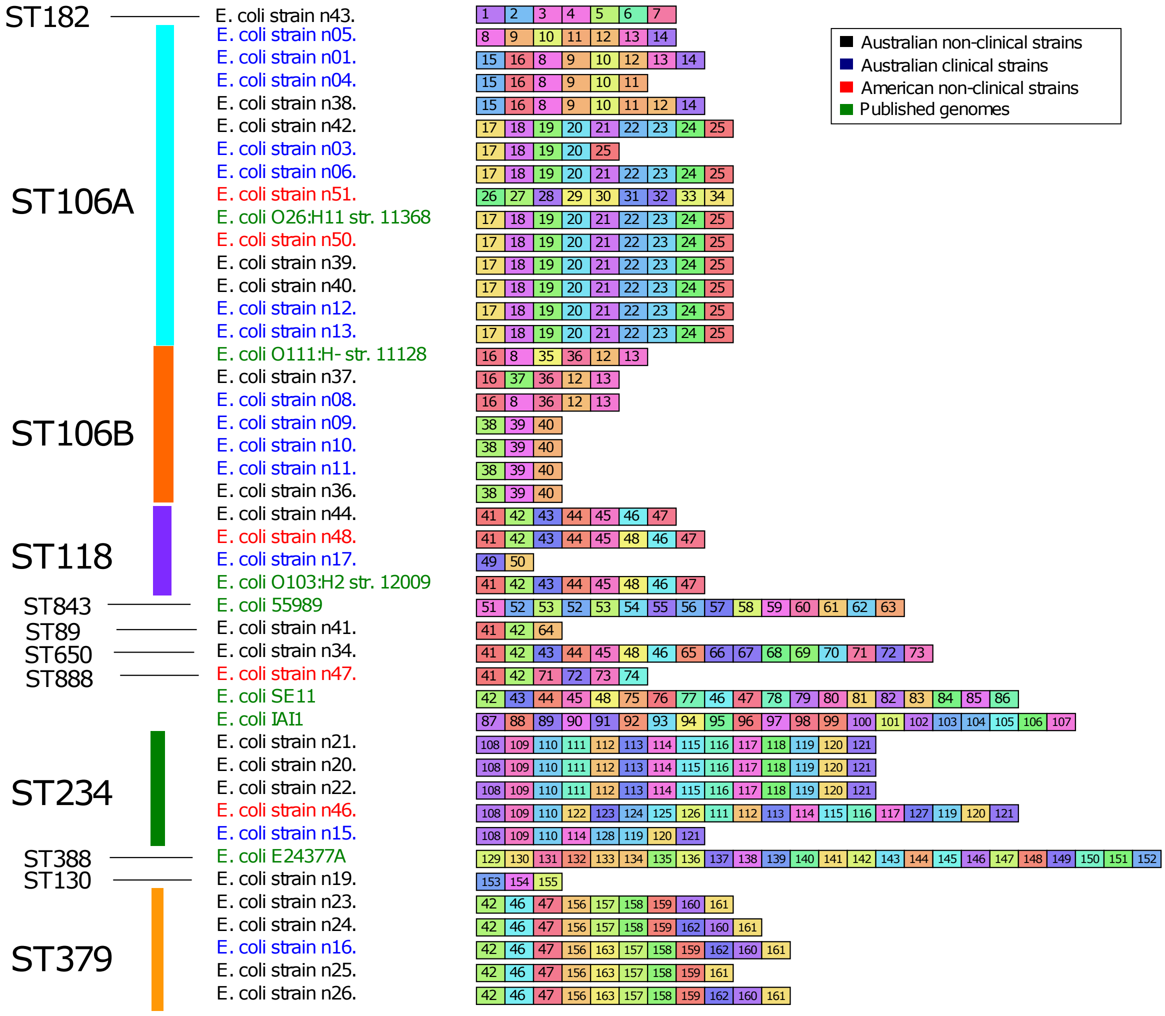

**Supplementary Figure 6.** Conservation of CRISPR2 spacers in non-O157 STEC and B1 phylogroup *E. coli*

Graphic representation of spacer content from CRISPR2 for non-O157 STEC strains and B1 phylogroup *E. coli* that share space sequences. A uniquely colored box and symbol combination designates each spacer sequence. The colors and the numbers were assigned arbitrarily and are different from those in Figure 4. Sequences are listed (left to right) from farthest to nearest the CRISPR leader sequence. Strains and lineages are listed in order and colored according to the scheme used for the phylogram in Figure 1.

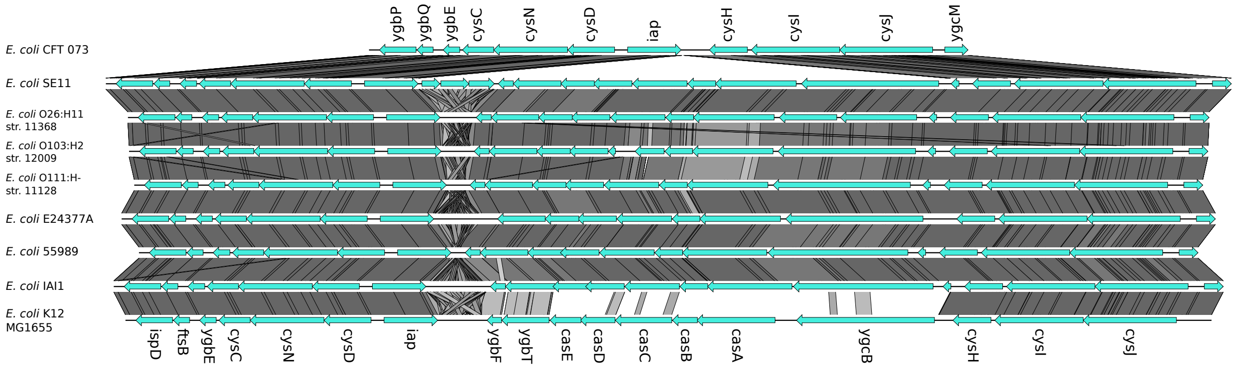

**Supplementary Figure 7.** tBLASTx comparison of CRISPR1 loci between *iap-cysH*

Translated nucleotide comparison (BLAST 2.2.27+ tBLASTx) [2] of *E. coli* CRISPR1 loci, between *iap* and *cysH*. The comparison included CFT073, K12MG1665, published non-O157 STEC strains (11368, 11128 and 12009) and other *E. coli* within the B1 phylogroup; SE11, ETEC E24377A, IAI1, EAEC 55989. Accession numbers for strains are listed in Table XXX. The comparison shows that *cas* genes for STEC and B1 strains were highly divergent to K12 and the region was absent in CFT073. Figure was prepared using EasyFig [5].

**Supplementary Table 1.** Common Shiga toxin variants

| **Gene name** | **Description** | **Source organism** | **Accession No.** |
|---|---|---|---|
| stx1 | Shiga toxin 1 | S.dysenteriae | M19437 |
| stx1 | Shiga toxin 1 | O157:H7 str. Sakai | BAB36396.1, BAB36397.1 |
| $stx1_{OX3}$ | Shiga toxin 1 (OX3) | O128:H2 | AJ413986 |
| stx1c | Shiga toxin 1c | *E. coli* phage 6220 | CAC88707 |
| stx2a | Shiga toxin 2a | O157:H7 str. Sakai | BAB34628, BAB34629 |
| stx2b | Shiga toxin 2b | ONT:H4 | AM498375 |
| stx2c | Shiga toxin 2c | O177:H | FM177472 |
| stx2d | Shiga toxin 2d | O22:H8 | HQ585061, HQ585062 |
| $stx2d_{activatable}$ | Shiga toxin AB 2d (mucus-activatable) | *E. coli* | CCA61220 |
| stx2e | Shiga toxin 2e | ONT:H- | AJ567998 |
| stx2f | Shiga toxin 2f | O128:H2 | AJ270998 |
| stx2g | Shiga toxin 2g | O2:H25 | AJ966782 |

**Supplementary Table 2.** Representative *E. coli* genomes

| **Strain name** | **Serotype** | **Accession** | **Pathotype** |
|---|---|---|---|
| 11368 | O26:H11 | AP010953 | EHEC |
| Sakai | O157:H7 | BA000007 | EHEC |
| 12009 | O103:H2 | AP010958 | EHEC |
| 11128 | O111:H- | AP010960 | EHEC |
| K12 MG1665 | | U00096 | Lab-adapted |
| IAI1 | | CU928160 | Commensal |
| HS | | CP000802 | Commensal |
| BL21 | | CP001665 | Lab-adapted |
| E24377A | | CP000800 | ETEC |
| 55989 | | CU928145 | EAEC |
| CFT073 | | AE014075 | UPEC |
| SE11 | | AP009240 | Commensal |

**Supplementary Table 3.** Common STEC virulence factors

| Gene name | Description | Source | Group | Accession no. | Start – Stop |
|---|---|---|---|---|---|
| LT-A | heat-labile enterotoxin subunit A | Plasmid EWD 299 | ETEC | M57244.1 | 1-723 |
| EibA | immunoglobulin binding protein | *E. coli* Eda1 | Fimbrae | CU928162.2 | 2441310-2440120 |
| EibG | immunoglobulin binding protein | *E. coli* str. 393/98 | Fimbrae | GU295813.1 | 1-1527 |
| elfA | E coli laminin binding fimbrae | O157:H7 str. EDL933 | Fimbrae | AE005174 | 1216554-1217102 |
| F9 | F9 fimbraie | O157:H7 str. EDL933 | Fimbrae | AE005174 | 1974046-1974609 |
| HcpA | haemorrhagic pilus | O157:H7 str. EDL933 | Fimbrae | AE005174 | 122039-121599 |
| lpfA | long polar fimbriae | O157:H7 str. EDL933 | Fimbrae | AAG58695.1 | 1-178 |
| SfaA | S-fimbrillin | *E. coli* O6:K15:H31 | Fimbrae | AAD02650.1 | 1-176 |
| FyuA | Ferric Yersiniabactin uptake receptor | *E. coli* UTI89 | HPI | CP000243 | 2098302-2100323 |
| Efa-1 | EHEC factor for adherence | O157:H7 str. EDL933 | O-Island 122 | AE005174 | 3934514-3935815 |
| Efa-2 | EHEC factor for adherence | O157:H7 str. EDL933 | O-Island 122 | AE005174 | 3935821-3936648 |
| nleB | non-LEE encoded effectors | O157:H7 str. EDL933 | O-Island 122 | AE005174 | 3930927-3931916 |
| nleE | non-LEE encoded effectors | O157:H7 str. EDL933 | O-Island 122 | AE005174 | 3931965-3932639 |
| pagC | phoP-activated gene C | O157:H7 str. EDL933 | O-Island 122 | AE005174 | 3925674-3925141 |
| sen | Shigella flexneri enterotoxin homolog | O157:H7 str. EDL933 | O-Island 122 | AE005174 | 3928670-3930319 |
| ureD | Urease accessory protein D | O157:H7 str. EDL933 | O-Island 43 | AE005174 | 1078653-1079351 |
| ureA | urease structural subunit A | O157:H7 str. EDL933 | O-Island 43 | AE005174 | 1079442-1079744 |
| ureB | urease structural subunit B | O157:H7 str. EDL933 | O-Island 43 | AE005174 | 1079753-1080073 |
| ureC | urease structural subunit C | O157:H7 str. EDL933 | O-Island 43 | AE005174 | 1080063-1081769 |
| ureE | Urease accessory protein E | O157:H7 str. EDL933 | O-Island 43 | AE005174 | 1081779-1082243 |
| ureF | Urease accessory protein F | O157:H7 str. EDL933 | O-Island 43 | AE005174 | 1082244-1082918 |
| ureG | Urease accessory protein G | O157:H7 str. EDL933 | O-Island 43 | AE005174 | 1082930-1083547 |
| terZ | Tellurite resistance | O157:H7 str. EDL933 | O-Island 43 | AE005174 | 1100663-1101244 |
| terA | Tellurite resistance | O157:H7 str. EDL933 | O-Island 43 | AE005174 | 1101244-1102401 |
| terB | Tellurite resistance | O157:H7 str. EDL933 | O-Island 43 | AE005174 | 1102424-1102879 |
| terC | Tellurite resistance | O157:H7 str. EDL933 | O-Island 43 | AE005174 | 1102902-1103942 |
| terD | Tellurite resistance | O157:H7 str. EDL933 | O-Island 43 | AE005174 | 1103991-1104569 |
| terE | Tellurite resistance | O157:H7 str. EDL933 | O-Island 43 | AE005174 | 1104638-1105213 |
| terF | Tellurite resistance | O157:H7 str. EDL933 | O-Island 43 | AE005174 | 1105635-1105943 |
| Aida-1 | autotransporter adhesin involved in diffuse adherence | O157:H7 str. EDL933 | O-Island 43 | AE005174 | 1132148-1135165 |
| ehxA | EHEC haemolysin | O113:H21 | Plasmid | AY258503 | 117132-114136 |
| epeA | *E. coli* autotransporter protease | O113:H21 | Plasmid | AAL18821 | 1-1359 |
| espP | extracellular serine protease autotransporter | O113:H21 | Plasmid | AAZ76514 | 1-1300 |
| etpD | type II secretion pathway related protein | O157:H7 | Plasmid | AB011549 | 3675-5432 |
| HlyA-ehx | EHEC haemolysin | O157:H7 str. Sakai | Plasmid | BAA31774 | 1-998 |
| katP | catalase-peroxidase | O157:H7 str. Sakai | Plasmid | BAA31832 | 1-736 |
| pilQ | Pili translocation | O113:H21 | Plasmid | AY258503 | 10095-11603 |
| saa | STEC autoagglutinating adhesin | O113:H21 | Plasmid | AY258503 | 143552-145156 |
| Sab | Autotransporter; biofilm formation | O113:H21 | Plasmid | AY258503 | 123200-118905 |
| stcE | Adhesion metalloprotease | O157:H7 | Plasmid | AF074613 | 23016-25712 |
| subA | subtilase cytotoxin | O113:h21 | Plasmid | AAZ76524.1 | 1-347 |
| subB | subtilase cytotoxin | O113:H21 | Plasmid | AAZ76523.1 | 1-141 |
| AstA | enteroaggregative heat-stable toxin 1 (EAST1) | *E. coli* 55989 | Toxin | AAM88299.1 | 1-38 |
| cdtA | cytolethal distending toxin subunit A | APEC O1 | Toxin | ABJ00842 | 1-237 |
| cdtB | cytolethal distending toxin subunit B | APEC O1 | Toxin | ABJ00843 | 1-273 |
| cdtC | cytolethal distending toxin subunit C | APEC O1 | Toxin | ABJ00844 | 1-190 |
| stx1 | Shiga-like toxin 1 subunit AB | O157:H7 str. Sakai | Toxin | BAB36396.1 | 1-89 |
| stx2 | Shiga-like toxin 2a subunit AB | O157:H7 str. Sakai | Toxin | BAB34629 | 1-89 |
| toxB | Host selective toxin | O26:H11 | Toxin | BAH24012.1 | 1-3166 |
| papA | structural subunit of F13 P-pili of UPEC | E coli str. 132 | UPEC | EU156123.1 | 1-555 |
| sodC | Copper zinc superoxide dismutase | O157:H7 str. EDL933 | | AAG56635 | 1-173 |
| tpx | thioredoxin-dependent thiol peroxidase | O157:H7 str. Sakai | | BAB35326 | 1-168 |
